# Impact of Acetate and Optimized Nitrate Levels on Mixotrophic Growth and Protein Dynamics in *Chlorella Sorokiniana*

**DOI:** 10.1101/2024.09.04.611160

**Authors:** Sunni Chen, Ruiqi Wang, Youn Joong Kim, Emily Radican, Yu Lei, Yongku Cho, Zhenlei Xiao, Mingyu Qiao, Yangchao Luo

## Abstract

Microalgae are well-known for their role as sustainable bio-factories, offering a promising solution to the global food and nutrition crisis. To clarify the potential of *Chlorella sorokiniana* UTEX 1230 for food applications, particularly as an alternative protein source, the study employed a mixotrophic cultivation mode with sodium acetate (NaAc) as a cost-effective organic carbon (NaAc-C) source. Varying levels of NaAc-C and nitrate-sourced nitrogen were investigated, optimizing the effect of metabolic characteristics of the microalgal growth. The designed heterotrophic cultivation confirmed the ability of *C. sorokiniana* UTEX 1230 to grow on NaAc-C, and then the mixotrophic cultures, when supported by both NaAc-C and CO_2_, exhibited superior growth performance, achieving double the biomass concentration compared to the autotrophic control. The addition of nitrogen (750 mg/L NaNO₃) facilitated the thorough metabolism of NaAc-C and enhanced photosynthetic activity indicated by a 196% increase in pigment levels, which resulted in a maximum biomass concentration of 2.82 g/L in the 150 mM NaAc-C group. A detailed analysis of nitrogen and protein concentrations over time revealed that higher nitrogen availability led to greater protein accumulation which was then degraded to support essential life activities under nitrogen starvation. Therefore, it is suggested that supplementing nitrate on the 3^rd^ day and harvesting on the 4^th^ day could be strategically implemented to increase protein yield from 0.17 g/L/d to 0.34 g/L/d. These findings offer theoretical guidance for further refining this microalgal strain for use as an alternative protein.

## 1. Introduction

Dietary proteins are critical for providing essential amino acids, which cannot be synthesized de novo for the constitution of body proteins, highlighting their significance to animal and human health. In 2017, the world population of 7.3 billion necessitated an estimated 202 million tons of protein per annum, with an expected increase of 57% by 2050[1]. Ensuring an adequate supply of dietary protein is challenging but pivotal in any consideration of global food and nutrition security [2]. Microalgae are unicellular algae typically found in marine and freshwater, possessing unique advantages over the cultivation of other traditional foods, such as a short growth cycle, independence from herbicides or antibiotics, no constraints on land conditions, and high tolerance to various environments [3]. Of great importance is the superior photosynthetic efficiency of microalgae (up to 20%), enabling them to function as sustainable bio-factories yielding carbohydrates, lipids, proteins, pigments, vitamins, etc., via sequestering CO_2_ [4] and thus positioning them as a crucial resource for the future of the food and nutrition industries. Currently, microalgal protein is considered a viable substitute for human food or animal feed [5–7], offering a quality comparable to traditional pulses like lentils, beans, peas, and chickpeas [8].

Except for the autotrophic capacity harvesting light energy and CO_2_ for biomass accumulation, some strains of microalgae could also hold the ability to utilize organic carbons for growing in the absence of light, called heterotrophic cultivation. It is feasible to consider mixotrophic cultivation, which combines autotrophic and heterotrophic cultivation by simultaneously providing light and organic carbon, aiming to double biomass production [9]. Available research suggests that the two processes may positively affect each other, contributing to a synergistic effect [10]. Currently, closed photobioreactors are gaining popularity due to a more controllable growth condition environment thus reduced risk of contamination, compared to open raceways [11]. This is especially true for mixotrophic microalgae cultivation, where high productivity of high-value compounds can offset costs associated with facilities, leading to a more economically feasible production. However, another factor that should be considered for large scale production and application is the additional costs of organic carbons, accounting for 80% of production costs in the heterotrophic section [12].

The primary organic carbons used for growing mixotrophic microalgae are sugar-type substrates, such as glucose [13], typically derived from terrestrial crops and thus come at a high price. To address this, seeking cheaper organic substrates is essential, with waste-sourced options holding significant promise. Volatile fatty acids (VFAs) can be produced from solid and liquid wastes through anaerobic fermentation, microbial electrosynthesis, or electrochemical processes based on inorganic catalysts [14], representing a hopeful alternative carbon source for microalgae. As proven, the VFA mixture obtained after acidogenic fermentation of waste lignocellulosic biomass from brewers’ spent grain and wastewater from the potato processing industry can effectively support the growth of *Chlorella sorokiniana* [15, 16]. In addition to the lower cost of acetate (300– 450 USD per ton) compared to glucose (500 USD per ton) [17], VFAs are less preferred by bacteria, thereby reducing the risk of contamination [18], further enhancing their value as an organic carbon source for industrial microalgal production.

The cultivation conditions could be finely tuned, including temperature, light intensity, photoperiod, pH, CO_2_ supply and so on, for improving the microalgal biomass content and composition and, consequently, the quality of the final product [19], among which nutrient availability shows the most significant influence and is relatively easy and economical to control. Besides carbon, nitrogen plays a critical role in microalgal cultivation as a critical element and a key component in high-value bioproducts, including proteins, chlorophylls, and DNA. Generally, many microalgal strains shift their carbon storage patterns toward neutral lipids or carbohydrates under nitrogen limitation but conversely, proteins accumulate when nitrogen is sufficient [20]. *C. sorokiniana* is one of the strains identified as Generally Recognized as Safe (GRAS) by the United States Food and Drug Administration (FDA) [21] and has also been shown to be sufficiently robust for growth conditions that are unfavorable for other algal strains [22]. Lizzul, Lekuona-Amundarain, Purton and Campos [23] have demonstrated that *C. sorokiniana* UTEX 1230 can grow on sodium acetate (NaAc) sourced carbon (NaAc-C), but the impact of NaAc-C and the combined effects of nitrogen levels on cellular metabolic compositions have not been thoroughly explored. Therefore, this study aimed to fill this gap, providing a theoretical foundation for the subsequent industrial utilization of this microalgae (especially *C. sorokiniana*) as food or food additives.

## 2. Materials & Methods

### 2.1. Chemical materials

Bold’s Basal Medium (BBM) was purchased from PhytoTech Labs (Kansas, USA), sodium nitrate from Fisher Scientific (USA), and all other reagents, including 4-(2-hydroxyethyl)-1-piperazineethanesulfonic acid (HEPES) buffer, NaAc, and the HPLC grade solvents, from Sigma-Aldrich (USA).

### 2.2 Microalgal strain and culture conditions

*C. sorokiniana* UTEX 1230 was purchased from the Culture Collection of Algae at The University of Texas at Austin (UTEX) and maintained in BBM in a homemade culture system (**Fig. 1**) under the following conditions: room temperature, 5% CO_2_ at a rate of 0.5 vvm, continuous light with the intensity of 150 μmol/m^2^/s, and magnetic stirring at a rate of 200 rmp. The initial pH of the medium was adjusted to 6.8 ± 0.1 using 1M KOH, and the detailed medium composition can be found in **Table S1**.

**Fig. 1.**
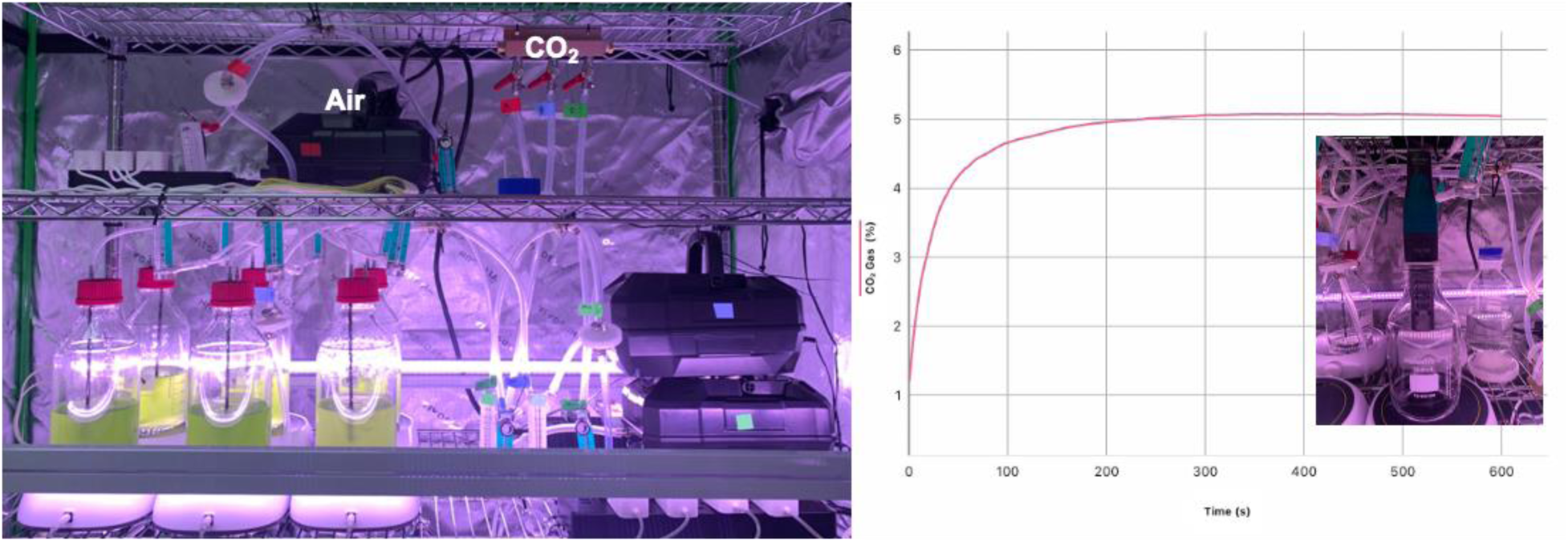
Schematic diagram of the culture system (Left), and the detection of CO_2_ level in the mixed gas via Vernier™ Go Direct™ CO_2_ Gas Sensor (Right).

All heterotrophic/mixotrophic flask experiments were initiated using the above autotrophic inoculum but supplemented with NaAc-C as the organic carbon source, while for heterotrophic cultivation, the bottles were covered with aluminum foil to block light. To ensure reliable data, cultures were adopted for at least one cultivation period following any change in conditions (see Sections 2.2 and 2.3), after which the pre-acclimated cultures were used as inoculum for subsequent experiments, which were started at an optical density (OD) at 750 nm of 0.1. During cultivation, 5 mL of suspension was collected daily for growth monitoring. After experiments, the samples were centrifuged at 8000 rpm for 10 min to obtain biomass sediment and supernatant, which were stored at -20°C until analysis. Considering the water loss due to gas release, 8 mL of autoclaved water was replenished daily. For pH maintenance, HEPES buffer was provided for autotrophic and heterotrophic cultivation.

### 2.3 Effects of sodium acetate concentrations as the organic carbon source

NaAc concentrations of 25, 50, 75, 100, and 125 mM, equivalent to 50, 100, 150, 200, and 250 mM NaAc-C, were studied for the mixotrophic cultivation of *C. sorokiniana* over 5 or 6 days. The growth rate was assessed as the response variable.

### 2.4 Effects of sodium nitrate concentrations as the nitrogen source

*C. sorokiniana* was cultivated with the addition of 100 or 150 mM NaAc-C in BBM (1N, 250 mg/L NaNO_3_) or in modified BBM with NaNO_3_ concentrations of 500 (2N), 750 (3N), 1000 (4N), 1250 (5N), and 1500 mg/L (6N). The growth rate and protein levels were evaluated as response variables.

### 2.5 Cell growth monitoring

The microalgal growth was monitored daily by OD changes at 750 nm using a spectrophotometer (Evolution 201, Thermo Scientific, USA). BBM, prepared prior to the start of batch fermentation, served as the blank in the analysis. The dry weight (DW) of the microalgae was determined through the following steps: i) Sampling the microalgae suspension; ii) Centrifuging at 8000 rpm for 10 min; iii) Washing twice with distilled water to remove excess nutrients; iv) Oven-drying at 50 °C until a constant weight was achieved. Because a good linear correlation fit was obtained (R^2^ value of 0.99) between the DW and OD measurements, the experiment applied the equation of the standard curve to determine the DW using the obtained OD value for convenience.

### 2.6 Kinetic growth parameters

The specific growth rate (*μ*, day^−1^) was calculated from Eq. (1), where *C*_1_ and *C*_2_ were the concentration of biomass (g/L) at the day *t*_1_ and *t*_2_ (d), respectively.

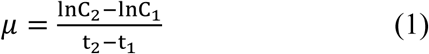

Biomass productivity (*P*, g/L/d) during the culture period was calculated from the Eq. (2)

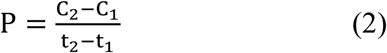

Accordingly, the highest levels could be assigned as the highest concentration of biomass *C_max_*, *μ_max_* and *P_max_*.

### 2.7 Nutrients remaining in microalgal cultures

#### 2.7.1 Acetate concentrations

The determination of acetate concentration was performed in the Vanquish™ HPLC system (Thermo Scientific, USA). Hypersil GOLD™ C18 column (4.6 ∼ 150 mm, 3 μm, Thermo Scientific, USA) with a temperature of 30 °C, and the mobile phase consisting of methanol, water and phosphoric acid at a ratio of 10/90/0.1 (v/v) was used to separate a 5 μL sample. The flow rate was held at 0.8 mL/min, and the total run time was 10 min. The signal was monitored at 230 nm by a variable wavelength UV−vis detector and quantified based on the external standard method.

#### 2.7.2 Nitrate concentrations

Referring to the assay reported by Witthohn, Schmidt, Strieth, Ulber and Muffler [24] with slight modification, the remaining NO_3_^-^ levels in the culture supernatant were determined using the sodium salicylate assay. Specifically, 25 μL of the sample was seeded into a 96-well microtiter plate, followed by the addition of 25 μL of 1% (w/v) sodium salicylate solution. The plate was then heated at 80°C for 1 hour in a dehydrator (Vevor, Home Depot, USA) to evaporate the water. Subsequently, the remaining salts were dissolved in 25 μL of concentrated H_2_SO_4_ by mixing on a shaker (Solaris, Thermo Scientific, USA) at 200 rpm for 20 min at room temperature. The acidic solution was then diluted with 150 μL of H_2_O and alkalized by adding 175 μL of 30% (w/v) NaOH. After an incubation period of 5 min at room temperature, the absorption was measured at 412 nm using a microtiter plate reader (Synergy H1, BioTek, USA). Quantification was achieved by establishing a standard curve using a series of sodium nitrate solutions.

### 2.8 Composition of biomass

#### 2.8.1 Protein content

The total protein content in the microalgae was determined by the Dumas method (AOAC 990.03), using an Elementar CHNS analyzer (vario MICRO cube, Germany), with a conversion factor of 6.25 [25].

#### 2.8.2 Lipid

The lipid was extracted via the Bligh and Dyer method with ultrasound assistance [26]. A 100 mg sample of biomass was mixed with 5 mL water, 5 mL methanol, and 2.5 mL chloroform, and then homogenized in a sonicator (Q SONICA, Q125, USA.) with a power rating of 125 W for 10 min in pulse mode (45 s on; 15 s off) at 50% amplitude. Supplied with the 2.5 mL chloroform, the mixture was subjected to the same ultrasonic condition for another 5 min. After centrifugation at 10,000 rpm for 10 min, the bottom chloroform layer containing lipids was collected in pre-weighed glass bottles. The total lipid content was then gravimetrically measured after solvent evaporation using an N-EVAP™111 nitrogen evaporator.

#### 2.8.3 Ash

The ash content was measured gravimetrically using thermalgravimetric analysis (TGA, TA instruments Q500-0188, USA) [27]. Around 10 mg of the sample was heated at a rate of 10 °C/min, held at 105 °C for 30 min, and then continued heating until reaching 800 °C for 30 min under an air atmosphere.

#### 2.8.4 Carbohydrate

The carbohydrate content of microalgae was calculated by deducting the percentages of protein, lipid, and ash from 100%.

#### 2.8.5 Chlorophyll a and b

The chlorophyll pigment content was measured referring to the method reported by Poad and Derek [28] with minimal modification. Briefly, the pellet from centrifugation was then re-suspended in 10 mL of 90% methanol solution and incubated overnight at 4 °C. Re-centrifugation of the extracts was then carried out at 8000 rpm for 10 min, and the absorbance of the resulting supernatant was read at 750 nm, 665 nm, and 660 nm. The following equations were then employed to calculate the pigment contents:

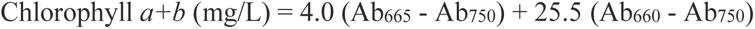

Where ‘Ab’ stands for absorbance at the specific wavelength. The absorbance results were subtracted from the absorbance at 750 nm for turbidity.

### 2.9 Data Analysis

Data was analyzed and plotted on Windows Microsoft Excel 2024 (Microsoft, London, UK).

Triplicate experimental results display error bars with standard deviations from the mean.

## 3. Result and Discussion

### 3.1 Determination of Initial Experimental Conditions

Mixotrophic cultivation has been demonstrated as a promising strategy to enhance microalgal biomass production by enabling the utilization of both organic and inorganic carbon sources under the presence of light at the same time. However, not all microalgal strains can effectively utilize organic carbon, nor is every organic carbon source equally suitable, meaning it is essential to confirm the potential heterotrophic capacity of microalgae with the desired carbon source, NaAc-C, in the present study. Although Lizzul, Lekuona-Amundarain, Purton and Campos [23] have already grown *C. sorokiniana* UTEX 1230 using NaAc-C, a more reliable and scientifically rigorous approach was adopted here to highlight the capability of utilizing NaAc-C by the microalgae. Specifically, heterotrophic cultivation was chosen to eliminate any contribution from the autotrophic section thus ensuring all biomass was derived solely from acetate or glucose. Additionally, to support the metabolism of organic carbon via heterotrophic cellular respiration, oxygen was supplied by introducing air into the medium. Besides, glucose, which is proven to support the optimal growth of chlorella [29], was used as the reference for comparison. Obviously, when cultured with glucose, the microalgae entered the exponential growth phase on the 5^th^ day, concluded on the 8^th^ day due to glucose depletion, as evidenced by the reduced sugar method. When supplied with NaAc-C, the growth curve exhibited a similar shape, although with a longer lag phase (**Fig. 2a, 2b**). This observation suggested that acetate held promise as an organic carbon source for the studied microalgae. Indeed, the previous screening results demonstrated that the promotive effect of acetate was inferior to glucose but superior to fructose, sucrose, lactose, glycerol, and citric acid (data not shown).

**Fig. 2.**
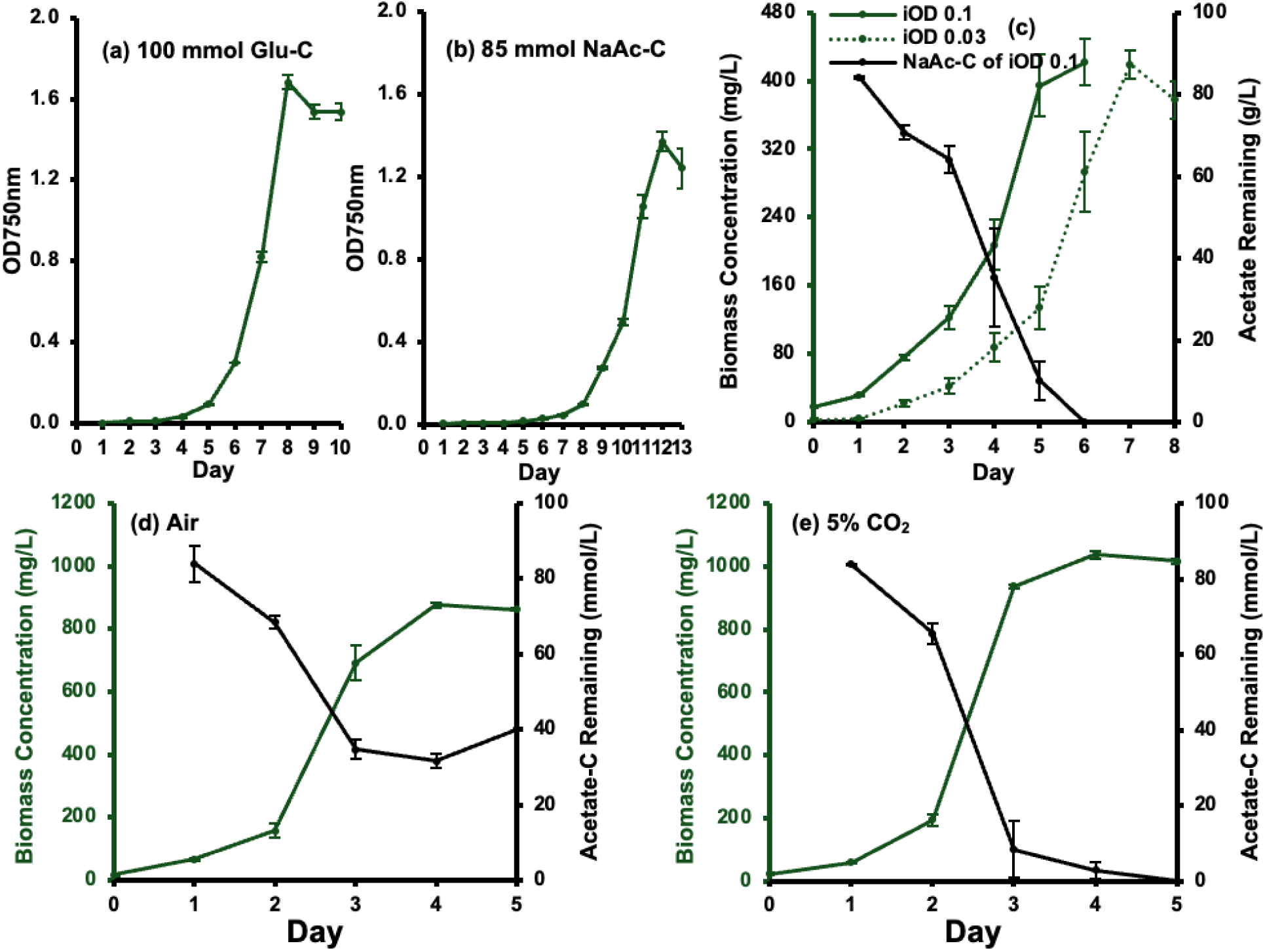
Growth curves of (a) Heterotrophic microalgae with glucose as the organic carbon source; (b) Heterotrophic microalgae with sodium acetate sourced carbon (NaAc-C); (c) Heterotrophic microalgae with NaAc-C at different inoculum sizes, including NaAc-C consumption (secondary axis in black); (d) Mixotrophic microalgae with NaAc-C and air, including NaAc-C consumption (secondary axis in black); (e) Mixotrophic microalgae with NaAc-C and 5% CO_2_, including NaAc-C consumption (secondary axis in black).

Shortening the lag phase is one of the primary considerations in microalgal large-scale culture. Typically, the inoculum size significantly impacts the kinetic growth parameters, including the lag phase, μ_max_, final biomass accumulation, and metabolite production [30]. **Fig. 2c** depicts the growth curve of heterotrophic microalgae with the initial optical density (iOD) of 0.03 (3 * 10^5^ cells/mL) and 0.1 (9 * 10^5^ cells/mL). It can be observed that the growth curves exhibited a similar shape, with similar kinetic growth parameters (**Table 1**), consistent with the curves between the 9% and 12% inoculum size drawn in the Cao, Kang, Gao, Wang, Pan and Liu [31]. But notably, a larger inoculum size resulted in a shorter lag phase due to a better acclimation ability to the new culture condition. A higher initial growth rate was due to an increase in divided cells, and an earlier entry into the stationary phase as a result of more rapid consumption of nutrients, which contributed to a shorter harvest time of 6 days [32]. Finally, the complete consumption of NaAc-C, as evidenced by HPLC analysis (**Fig. 2c**; **Fig. S1**, HPLC chromatogram), limited the ultimate biomass concentration.

**Table 1.** Kinetic growth parameters when sodium acetate as the organic carbon source.

|  | (c) iOD 0.03 | (c) iOD 0.1 | (d) Air | (e) 5% CO<sub>2</sub> |
| --- | --- | --- | --- | --- |
| $\mu_{\max}$ (/d) | 0.71 ±<br>0.01 | 0.71 ±<br>0.01 | 1.18 ±<br>0.04 | 1.24 ±<br>0.03 |
| $C_{\max}$ (mg<sub>dw</sub>/L) | 419.60 ±<br>16.58 | 421.84 ±<br>27.63 | 875.20 ±<br>8.96 | 1037.83 ±<br>13.70 |
| $P_{\max}$ (mg<sub>dw</sub>/L/d) | 146.76 ±<br>28.42 | 171.12 ±<br>9.27 | 533.76 ±<br>33.05 | 743.22 ±<br>13.11 |

\*c, d, e referred to the corresponding growth curve in Figure 2.

Mixotrophic cultivation was initiated using the same conditions of 85 mM NaAc-C, an 0.1 iOD, and aeration, with the additional removal of the aluminum foil covering from the bottle to allow light exposure. The results in **Table 1** confirmed the advantage of the mixotrophic cultivation, where the growth achieved 1.5 times the μ_max_, 2 times of C_max_, and 3 times of P_max_ as much as those in the heterotrophic cultivation (1.18 vs. 0.78 d^-1^; 875.20 vs. 421.84 mg_dw_/L; 533.76 vs. 171.12 mg_dw_/L/d, respectively). Ambient air usually serves as an inorganic carbon source, especially in large-scale industrial production [33], while it is well recognized that the supplementary addition of CO_2_ promotes biomass accumulation further for most microalgae [34]. To emphasize the importance of CO_2_ for the current strain, a comparative study on the mixotrophic growth of microalgae was conducted, where they were cultivated in a medium aerated with pure air versus 5% CO_2_-enriched air. The tolerance of microalgae to CO_2_ levels varies among strains, but 5% CO_2_ is generally considered safe for most strains [35]. Similarly, when provided with CO_2_, the microalgae exhibit greater μ_max_, C_max_, and P_max_, compared to those offered with air alone (**Table 1**). As indicated by the level of remaining NaAc-C in the medium (**Fig. 2c, d, and e**), the lower biomass production in the pure air group was partly due to the impaired capacity of the microalgae to utilize NaAc-C. The damage was inferred from the increased pH during the metabolic conversion of acetate to NaOH, where, without HEPES buffering, the pH reached up to 9.85 before the exponential phase, significantly exceeding the suitable pH for *C. sorokiniana* [36, 37]. Additionally, the absorption of acetate was reduced at a high pH where the proton transporters for transportation were limited [38]. In contrast, under the adjustment via 5% CO_2_, the pH remained 7 ∼ 8 throughout the entire period, supporting the well microalgal growth. The pH stabilization has been highlighted as essential for the optimal growth of *C. sorokiniana* on acetate [39] and for investigating the influence of other culture parameters, such as nutrients in the media [40].

Bicarbonate (HCO₃⁻), an ionic form of inorganic carbon in aqueous media, has recently gained interest as a potential carbon source for microalgal cultivation. HCO₃⁻ can help reduce the costs associated with CO₂ gas transportation and, more importantly, increase the availability of inorganic carbon [41]. As confirmed, only dissolved CO_2_ in the liquid phase could be utilized by microalgae for growth. In other words, the dissolution process becomes a critical bottleneck in the rate of CO_2_ fixation by microalgae [42]. However, concerns also arise from the high initial pH and the increased pH resulting from NaOH production after CO₂ absorption [43]. Given these considerations, it is recommended to use HCO₃⁻ for microalgae strains that favor high pH, such as *Spirulina*.

In summary, the microalgae were initially cultivated from an OD of 0.1 with a continuous supply of 5% CO_2_ to establish a stable growth baseline. Subsequently, the detailed effects of NaAc-C on microalgal growth were explored, which involved examining various concentrations of NaAc-C to assess their impact on growth rate, biomass accumulation, and overall metabolic responses.

### 3.2 Effect of Acetate Concentration

The effect of NaAc-C concentration was investigated under the previously established initial conditions. The biomass level almost reached its maximum on the third day when supplied with 50 mM NaAc-C, while on the same day, the biomass in the autotrophic control group was less than half (**Fig. 3a**), indicating that the accumulation was the combined result of autotrophy and heterotrophy. Interestingly, the OD values monitored at 750 nm did not show the same disparity (0.88 vs. 1.28). A similar discrepancy between cell density and dried biomass weight was also found by Miyawaki, Mariano, Vargas, Balmant, Defrancheschi, Corrêa, Santos, Selesu, Ordonez and Kava [44], which could be attributed to variations in cellular biochemical composition under different cultivation conditions. This suggested the necessity of re-establishing the OD value-biomass weight curve whenever cultivation conditions changed. Given a longer cultivation time (day 4∼6), the biomass did not significantly accumulate further due to the deficiencies of nutrients, including the depletion of NaAc-C (**Fig. 3b**), and the self-shading effect for the autotrophic part. Self-shading refers to the light attenuation by microalgae cell absorption that reduces light availability in the inner parts of the photobioreactors, and is exacerbated when the suspension has a high density or dark color [45], partly explaining the slower growth rate observed in the autotrophic group during the later phase (**Fig. 3a**)

**Fig. 3.**
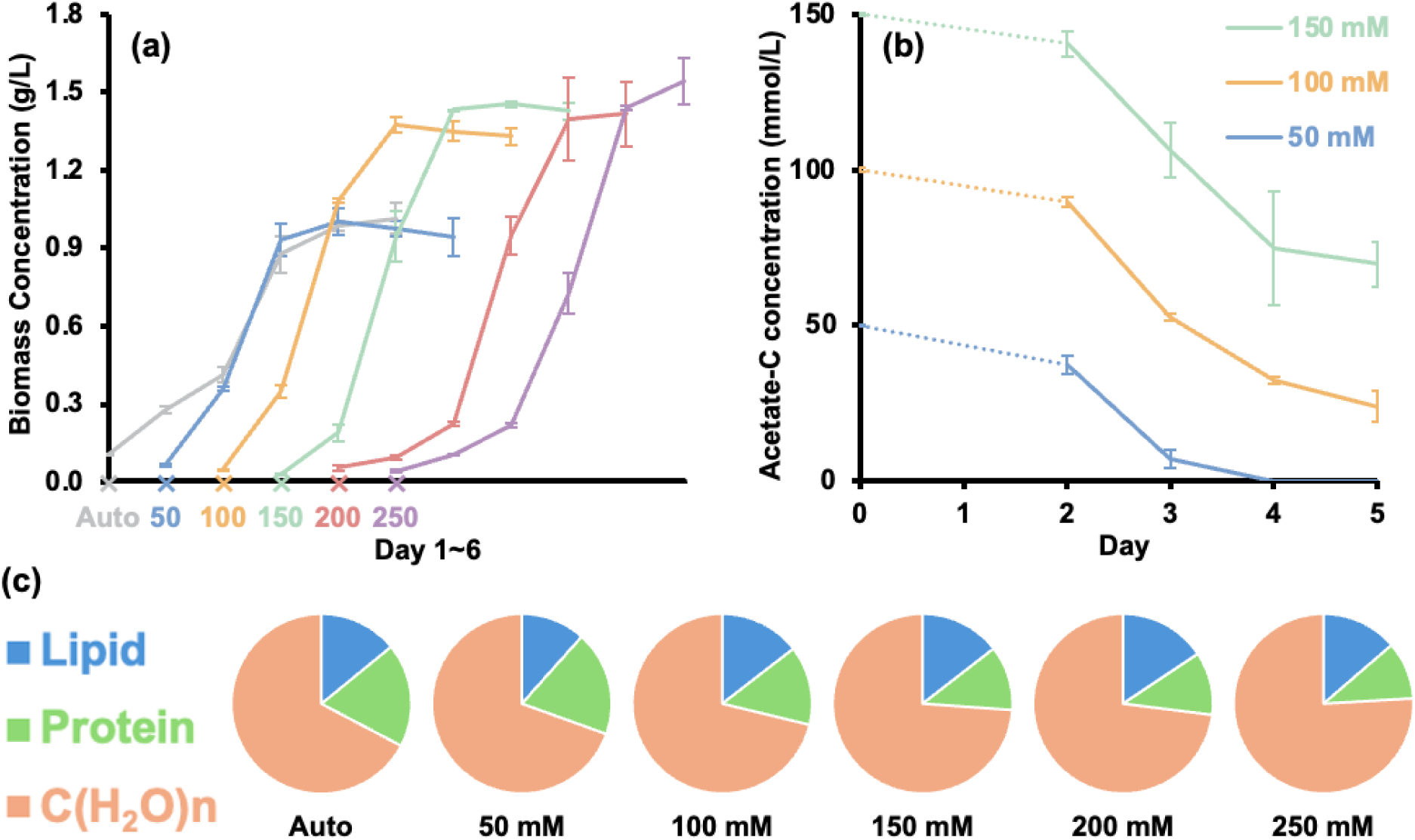
Effect of sodium acetate sourced carbon (NaAc-C) concentration on (a) growth curve; (b) NaAc-C consumption; (c) biochemical composition of microalgae. For clarity, the growth curve (a) was drawn separately, where the label below the x-axis indicated the treatment groups from autotrophy with 0mM NaAc-C to mixotrophy with 50 ∼ 250 mM NaAc-C, and each node on the curve represents a day over the entire 6-day period. The dotted line of (b) means not measured time point.

It was evident that higher NaAc-C concentrations resulted in prolonged adaptation times. Specifically, the lag phase was 0 days in the 0 NaAc-C groups, 1 day in the 50 and 100 mM NaAc-C group, 2 days in the 150 mM NaAc-C group, and 3 days in the 200 and 250 mM NaAc-C groups (**Fig. 3a**). Therefore, although higher biomass was expected over time, cultivating microalgae with more than 200 mM NaAc-C was not effective; while the treatment of 100 mM and 150 mM NaAc-C endowed the growth great, even if not the best, μ_max_, C_max_, and P_max_ (**Table 2**). The presence of residual NaAc-C (**Fig. 3b**) indicated that the growth of *C. sorokiniana* UTEX 1230 was likely limited by the insufficiency of other essential nutrients, underscoring the critical need to optimize the medium composition. The effect of pH on the increased lag time took no responsibility, as the initial pH change after the addition of NaAc-C was negligible, ranging from 6.89 in the 50 mM NaAc-C group to 7.02 in the 250 mM NaAc-C group. Instead, salinity was more likely the contributing factor, as exhibited in a study that explored the effect of different salinity levels on the growth performance of *Chlorella vulgaris.* This microalgae strain experienced a lag phase until Day 2 for 24 ppt and underwent a lag phase until Day 3 for 30 ppt [46]. A higher amount of NaAc-C did not always support higher biomass production; in contrast, a reduction of biomass accumulation was found when microalgae were fed 300 mM NaAc-C, and when treated with 400 and 800 mM NaAc-C no growth was observed within the first 6 days (**Fig. S2**). The substrate inhibition might be involved here, a common phenomenon when microalgae grow on VFAs [39]. The main cause of such inhibition is the accumulation of undissociated acid (CH_3_COOH) in a lower pH. This undissociated acid is lipophilic, allowing it to cross the membrane and enter cells, where it dissociates in the intracellular neutral environment, leading to cytosol acidification and anion accumulation [47]. Although the initial culture pH was neutral for the 300, 400, and 800 mM groups, the rapid introduction of CO_2_ and the ineffective metabolism of NaAc-C led to a pH decrease, producing a significant amount of CH_3_COOH and finally leading to substrate inhibition.

**Table 2.** Kinetic growth parameters treated with different NaAc-C levels.

|  | Auto | 50 mM | 100 mM | 150 mM | 200 mM | 250 mM |
| --- | --- | --- | --- | --- | --- | --- |
| **$\mu_{\max}$ (/d)** | 0.68 ± 0.03 | 1.30 ± 0.02 | 1.54 ± 0.01 | 1.69 ± 0.11 | 1.14 ± 0.08 | 0.98 ± 0.06 |
| **$C_{\max}$ (g<sub>dw</sub>/L)** | 1.02 ± 0.06 | 0.97 ± 0.03 | 1.37 ± 0.03 | 1.45 ± 0.01 | 1.42 ± 0.12 | 1.54 ± 0.09 |
| **$P_{\max}$ (g<sub>dw</sub>/L/d)** | 0.46 ± 0.04 | 0.57 ± 0.06 | 0.74 ± 0.03 | 0.76 ± 0.06 | 0.72 ± 0.07 | 0.75 ± 0.06 |

The biochemical compositions of microalgal biomass under different NaAc-C levels were investigated. As shown in **Fig. 3c**, one of the notable features was the low lipid percentage in the microalgae, consistent with the findings of a similar study [48]. The relatively higher lipid level in autotrophic cultivation (13.44%) compared to mixotrophic cultivation (11.09% for 50 mM NaAc-C) was similarly observed by Kobayashi, Noel, Barnes, Watson, Rosenberg, Erickson and Oyler [49] and Chai, Shi, Huang, Guo, Wei, Guo, Li, Dou, Liu and Liu [13], and the latter study also agreed that increased organic carbon promoted lipid accumulation (around 13% for 100 ∼ 250 mM NaAc-C). These results confirmed that the microalgae naturally produce less lipid regardless of the cultivation conditions, meaning that *C. sorokiniana* UTEX 1230 may not be well suited for biodiesel production, especially when compared to other strains with lipid content of up to 70% [50]. Interestingly, the high carbohydrate level in the used microalgae (up to 70% in 250 mM NaAc-C) enabled them to be great feedstock for biofuel production, including bioethanol, biobutanol, biomethane, and biohydrogen [51].

Mixotrophic *C. sorokiniana* UTEX 1230 exhibited a low protein concentration due to nitrogen deficiency (**Fig. 3c & Fig. S3**), a critical element for protein synthesis. There were only 13.66% and 11.15% in the 100 mM NaAc-C and 150 mM NaAc-C groups, respectively. During nitrogen starvation, the microalgae primarily metabolized organic carbon to synthesize carbohydrates, as indicated by the increased carbohydrate levels with rising NaAc-C concentrations (**Fig. 3c**). Correspondingly, proteins were degraded to generate carbon metabolic intermediates for biosynthesis. These metabolic alterations are consistent with the findings of Xiao, Zhou, Liang, Lin, Zheng, Chen and He [52], who also presented the potential of converting carbohydrates to protein via tuning the feed of carbon and nitrogen sources, inspiring the present study. Moreover, the intrinsic characteristic of *C. sorokiniana* UTEX 1230, characterized by low lipid production, left around 90% space for such adjustment, and importantly, this simplification of metabolic interactions primarily between carbohydrates and proteins, rather than involving lipids, allows for more straightforward optimization strategies.

In summary, 100 mM NaAc-C and 150 mM NaAc-C were identified as optimal carbon levels for microalgal growth, providing a faster growth rate and higher biomass accumulation. However, the remaining NaAc-C **(Fig. 3b)** and the unbalanced composition **(Fig. 3c)** indicated opportunities for further optimization, particularly concerning nitrogen supply.

### 3.3 Effect of Nitrate Concentration

#### 3.3.1 Effect of Nitrate Concentration on the Growth Curve

Microalgae can utilize various forms of nitrogen, including nitrate, nitrite, ammonium, and urea, depending on the strains, to support cell growth and biochemical synthesis. Among these, nitrate is more commonly used in microalgae cultivation due to its stability and lower likelihood of pH fluctuations in the culture medium compared to ammonium salts [53]. Therefore, additional NaNO_3_ was supplied as the nitrogen source to the medium to investigate its effect on microalgal growth, ranging from 500 mg/L (2N) to 1000 mg/L (5N) for the 100 mM NaAc-C group, while a higher supplementation did not contribute to increased biomass accumulation based on the preliminary experiment (**Fig. S4**). As depicted in **Fig. 4a**, due to a higher salinity brought by the addition of NaNO_3_, the lag phase in all groups was extended from 1 day to 2 days. Comparatively, all nitrogen-supplemented groups exhibited a higher C_max_ and P_max_, with the 3N group achieving the highest C_max_ of 2.38 g_dw_/L on the 5th day and displaying a one-fold increase in P_max_ (**Table 3**). The bell-shaped dependence of the biomass weight on the N level was also highlighted by relevant research [54, 55]. The increment can be attributed to the enhanced utilization of NaAc-C under balanced nutrient conditions (**Table 4**), as well as the greater availability of nitrogen (**Fig. 5**), while the decrease in biomass at higher nitrogen levels (4N and 5N) was likely caused by induced oxygen stress.

**Fig. 4.**
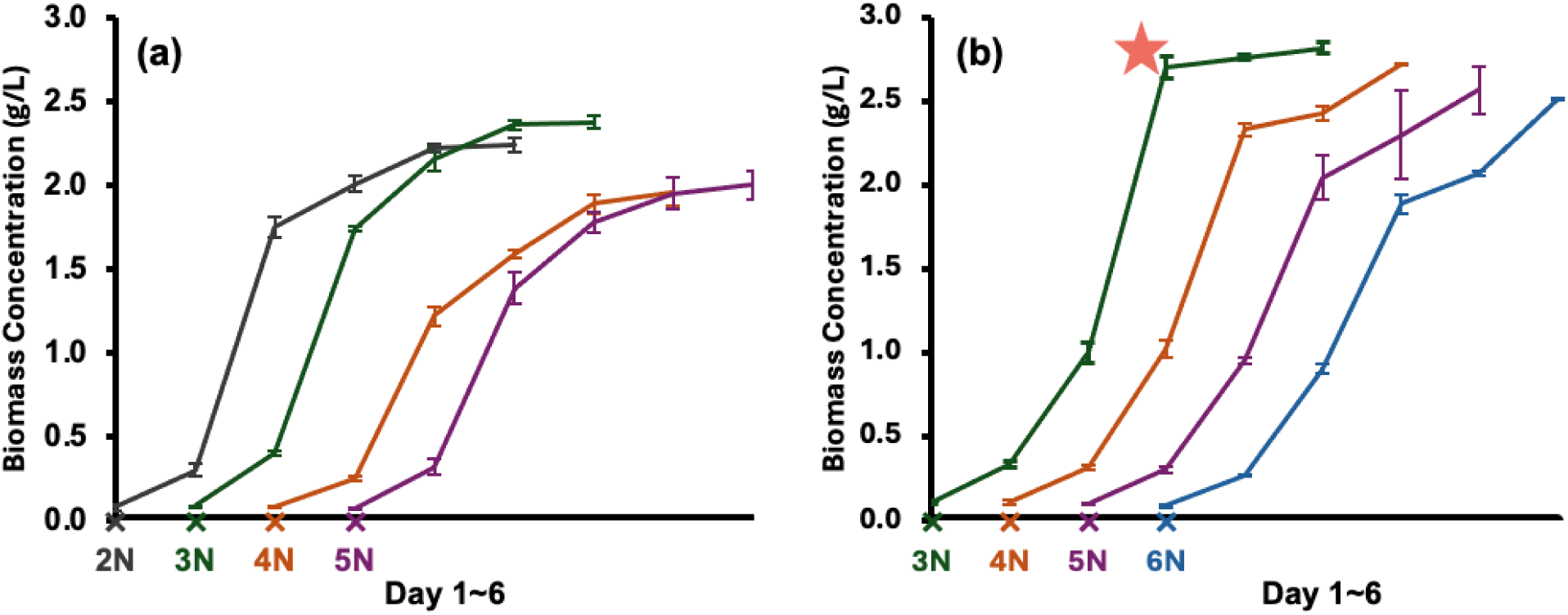
Effect of NaNO_3_ concentrations on (a) growth curve treated with 100 mM NaAc-C; (b) growth curve treated with 150 mM NaAc-C; (c) biochemical composition of microalgae. For clarity, the growth curve was drawn separately, where the label below the x-axis indicated the treatment groups from 2N (500 mg/L NaNO_3_) to 6N (1500 mg/L NaNO_3_), and each node on the curve represents a day over the entire 6-day period. **Star marked the recommended collection day or turning point for continuous cultivation**.

**Fig. 5.**
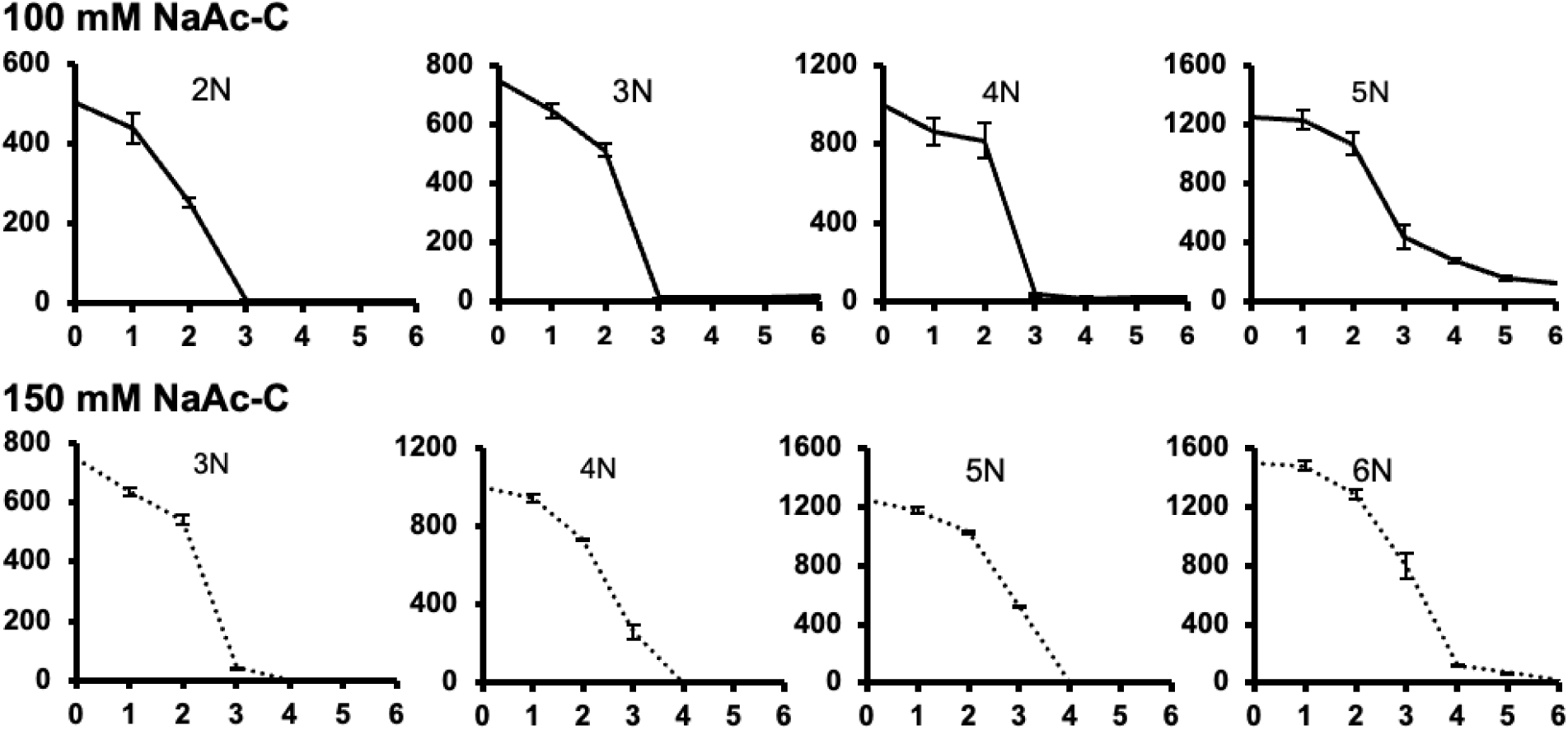
Effect of NaNO3 concentrations on (Up) NO_3_^-^ remaining in the medium when treated with 100 mM NaAc-C; (Down) NO_3_^-^ remaining in the medium when treated with 150 mM NaAc-C. The X-axis referred to cultivation time (d), and the Y-axis was the remaining NO_3_^-^ content (mg/L). 1N was 250 mg/L NaNO, 2N was 500 mg/L, and so forth.

**Table 3.** Kinetic growth parameters treated with different NaNO_3_ levels.

|  | 100 mM NaAc-C |  |  |  | 150 mM NaAc-C |  |  |  |
| --- | --- | --- | --- | --- | --- | --- | --- | --- |
|  | 2N | 3N | 4N | 5N | 3N | 4N | 5N | 6N |
| $\mu_{\max}$ (/d) | 1.55<br>± 0.11 | 1.49<br>± 0.01 | 1.33<br>± 0.02 | 1.45<br>± 0.04 | 1.08<br>± 0.01 | 1.04<br>± 0.06 | 1.02<br>± 0.04 | 1.04<br>± 0.03 |
| $C_{\max}$ (gdw/L) | 2.01<br>± 0.05 | 2.38<br>± 0.04 | 1.96<br>± 0.85 | 2.00<br>± 0.08 | 2.82<br>± 0.03 | 2.72<br>± 0.00 | 2.57<br>± 0.14 | 2.52<br>± 0.01 |
| $P_{\max}$ (gdw/L/d) | 1.45<br>± 0.07 | 1.34<br>± 0.03 | 0.97<br>± 0.05 | 1.06<br>± 0.07 | 1.70<br>± 0.00 | 1.31<br>± 0.02 | 1.09<br>± 0.11 | 0.98<br>± 0.03 |

1N was 250 mg/L NaNO<sub>3</sub>, 2N was 500 mg/L, and so forth.

**Table 4.** NaAc-C remaining treated with different NaNO_3_ levels.

|  | 100 mM |  |  |  | 150 mM |  |  |  |
| --- | --- | --- | --- | --- | --- | --- | --- | --- |
|  | 2N | 3N | 4N | 5N | 3N | 4N | 5N | 6N |
| 0 | 101.28<br>± 1.89 | 108.75<br>± 1.56 | 100.42<br>± 0.45 | 108.44<br>± 2.00 | 153.24<br>± 1.53 | 155.92<br>± 1.51 | 150.67<br>± 2.17 | 153.41<br>± 5.46 |
| 3 | 10.29<br>± 0.45 | ND | ND | ND | 85.18<br>± 4.06 | 85.32<br>± 11.27 | 89.78<br>± 0.59 | 96.80<br>± 12.03 |
| 4 | ND | ND | ND | ND | ND | ND | ND | 12.63<br>± 2.31 |
| 5 | ND | ND | ND | ND | ND | ND | ND | ND |
| 6 | ND | ND | ND | ND | ND | ND | ND | ND |

1N was 250 mg/L NaNO<sub>3</sub>, 2N was 500 mg/L, and so forth.
ND: Non-Detected

Combined with the treatment of 150 mM NaAc-C (**Fig. 4b**), the best microalgae growth performance was presented in the 3N group, even better than the 100 mM NaAc-C * 3N group, with the highest C_max_ of 2.82 g_dw_/L and P_max_ of 1.70 g_dw_/L/d, higher than those observed in most microalgal strains fed with NaAc-C [14]. The calculated C/N ratio was 14.45, falling within the well-recognized range of an optimal C/N ratio for microalgal growth. Although higher biomass production might be achieved over an extended period, the reduced growth rate under higher salinity, caused by the addition of nitrogen (**Fig. 4b**, 6N) and carbon (**Fig. S5**, 200 mM NaAc-C), rendered the cultivation less efficient. In fact, with the supplementation of 150 mM NaAc-C and 3N, 95% of the biomass accumulated within 4 days of cultivation, after which the depletion of carbon and nitrogen resources lowered the growth rate (**Fig. 5 and Table 2**). This indicated that fed-batch or continuous culture, with intermittent or constant nutrient supply, would be a more promising strategy for production, as demonstrated in other research [56]. Additionally, as previously discussed, higher cell density and/or a darker culture color lead to a stronger self-shading effect, necessitating increased light supply in the later phase, whereas such a strong light intensity should induce stress on the initial pale culture. Therefore, fed-batch light supplementation was also developed to compensate for the loss of light energy, resulting in higher biomass production or targeted bioproduct fields [57, 58]. These efforts have inspired the next step of the study, which will involve cultivating the microalgae in a photobioreactor in future studies.

#### 3.3.2 Effect of Nitrate Concentration on the Metabolic Composition

The images of microalgal samples visually revealed the changes in the green color of the microalgal cultures under varying nitrogen levels (**Fig. S6**), further warranting an investigation into the chlorophyll (a and b) content (**Fig. 6**). In the autotrophic cultivation, the nitrogen-containing chlorophyll was synthesized at the beginning when both nitrogen and light were available, but was then degraded for reutilization of the nitrogen so as to support further cell growth when the nitrogen was depleted for 3 days from the medium (**Fig. 6**, Auto * 1N) [59]. Mixotrophic cultivation reduced the photosynthetic activities, continuously sparing nitrogen for basic biomass accumulation rather than advanced pigment synthesis (**Fig. 6**, 100 mM NaAc-C * 1N and 150 mM NaAc-C * 1N), the same pattern observed in *Asterarcys sp.* SCS-1881 [60] and *Chlorella vulgaris* 31 [61]. With the addition of nitrogen, pigment changes over time followed a consistent pattern: acclimatization required about one day, after which higher nitrogen availability was associated with a greater increase in chlorophyll content until nitrogen depletion limited further biosynthesis (**Fig. 6**, middle and bottom rows), in accordance with the image evidence (**Fig. 6S**). Microalgae-derived pigments, including chlorophylls, carotenoids and phycobiliproteins, have unique colors and molecular structures, respectively, and show different physiological activities and health effects in the human body, thus gaining a high value in the global food pigment market [62]. The microalgae collected in the present study exhibited a significantly higher pigment level (1.35%, as observed in the 150 mM NaAc-C * 3N group) compared to the average 0.5 ∼ 1.0% chlorophyll content per DW [62], underscoring their potential as a natural pigment provider.

**Fig. 6.**
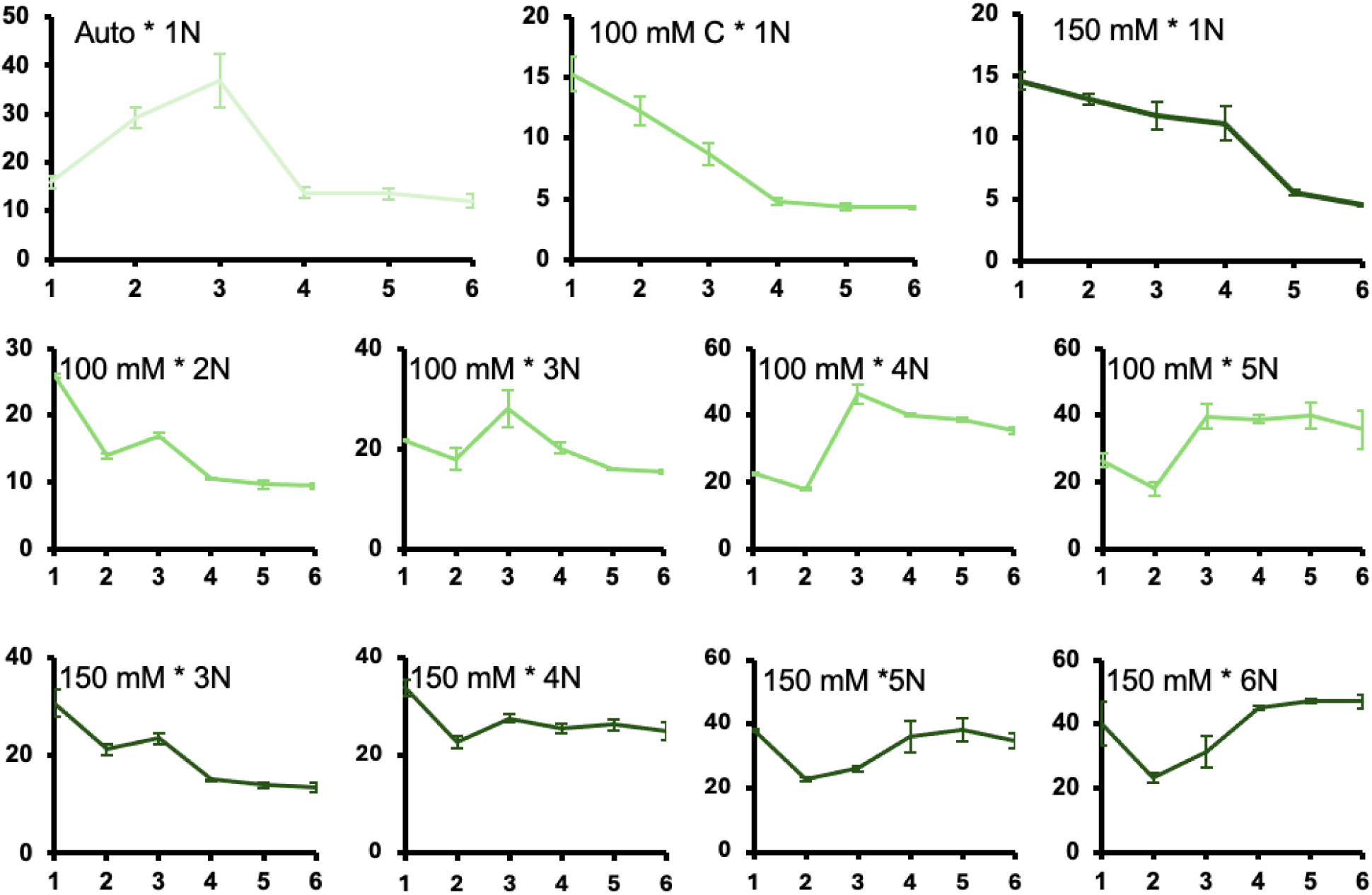
Changes of Pigment content in the autotrophic group (Light green), in the mixotrophic groups with the treatment of 100 mM NaAc-C and different nitrogen levels (Regular green) and in the mixotrophic groups with the treatment of 150 mM NaAc-C and different nitrogen levels (Dark green). The X-axis referred to cultivation time (d), and the Y-axis was pigment content (mg/g_dw_). 1N was 250 mg/L NaNO_3_, 2N was 500 mg/L, and so forth.

Due to difficulties in quantifying CO₂ gas consumption, it is challenging to determine the respective contributions of autotrophy and heterotrophy to mixotrophic biomass production based on the current results. An alternative approach involves supplying CO₂ in its ionic forms, such as HCO₃⁻ and CO₃²⁻, allowing for the calculation of biomass derived from inorganic carbon, whereas the consumption of organic carbon indicates heterotrophic biomass [63]. Briefly, this study suggested that the initial rapid biomass accumulation was driven by active photosynthesis and the high availability of organic carbon supporting respiratory processes. Subsequently, as organic carbon was depleted, slower biomass growth was primarily determined by the reduced photosynthesis rate, continuing until other essential nutrients were exhausted and growth entered a decline phase [45].

Although nitrogen levels are typically manipulated to enhance lipid and fatty acid production [64], the variations in nitrogen in this study were unable to overcome the inherent limitations of *C. sorokiniana* UTEX 1230 in lipid production, yet as the nitrogen supply increased, more proteins were synthesized in both the 100 and 150 mM NaAc-C groups (**Fig. 7**). The lower protein level in the 150 mM NaAc-C group compared to the 100 mM NaAc-C group under the same nitrogen treatment can be explained by the metabolic shift towards carbohydrate production due to the higher carbon input, consistent with the previous observation (**Fig. 3c**). Protein productivity, considering the biomass yield and cellular protein content together, is a more reliable index when protein is the target product. The 150 mM NaAc-C * 6N group had the highest yield (0.21 g/L/d) after 6 days of cultivation among all the C * N combinations. However, learning from the profound impact of nitrogen availability on protein metabolism, microalgal protein levels at different sampling times were also analyzed. Results shown in **Fig. 7b** indicated that when nitrogen was sufficient until the 3^rd^ day, microalgae uniformly accumulated around 50% protein across all 150 mM NaAc-C * N groups, ranking at the top [7]. Subsequently, the protein was degraded to support essential life activities, with the degree of degradation strongly dependent on the duration of nitrogen starvation. For the group of 150 mM NaAc-C * 3N, protein yield was calculated as 0.17 g/L/day for the first 3 days. As previously mentioned, fed-batch cultivation is an effective strategy to counteract the negative impact of high substrates on salinity to maximize the positive effect on growth. Therefore, considering the patterns of biomass accumulation **(Fig. 4b)**, protein changes **(Fig. 7b)**, and nitrate consumption **(Fig. 5)**, it is recommended to supply additional nitrate (250 mg/L) on the 3^rd^ day and then to collect samples on the 4^th^ day, ideally doubling the protein yield to 0.34 g/L/d (starred curve in **Fig. 4b**).

**Fig. 7.**
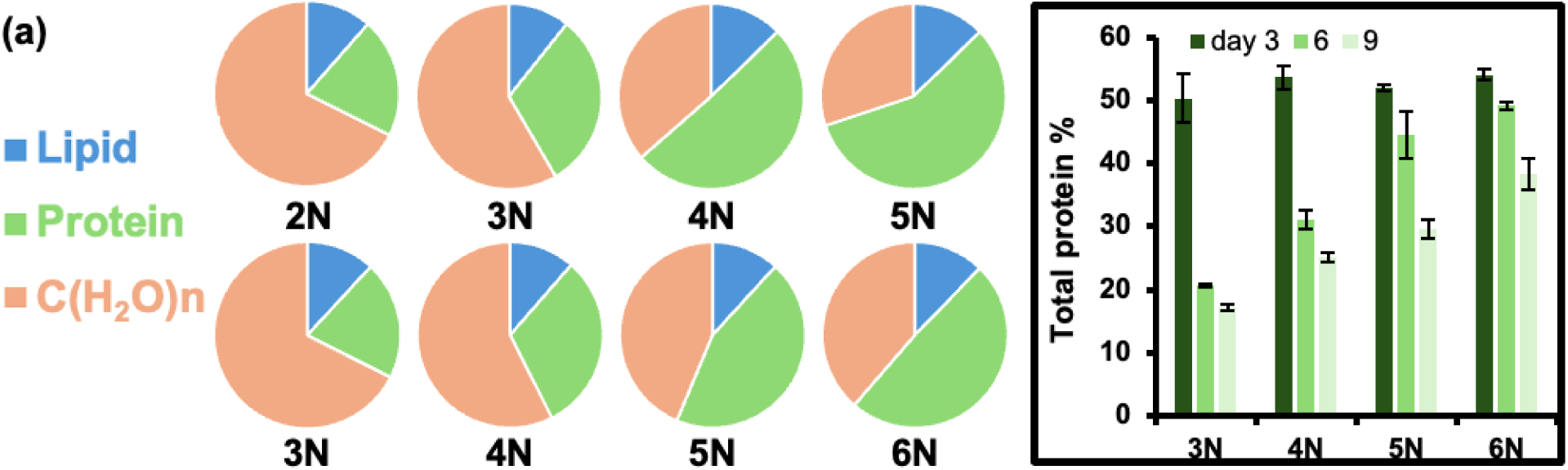
Effect of NaNO_3_ concentrations on (a) microalgal biochemical composition when treated with 100 mM NaAc-C (up) and microalgal biochemical composition when treated with 150 mM NaAc-C (Down) collected on Day 6; (b) microalgal protein level on different collection day composition when treated with 150 mM NaAc-C. 1N was 250 mg/L NaNO_3_, 2N was 500 mg/L, and so forth.

**Table 5** lists *C. sorokiniana* and other microalgal cultures using acetate as the carbon source, identified from Web of Science using the terms *((AB=(acetate)) AND AB=(microalga*)) AND AB=(protein)* reported over the past ten years. As shown, acetate could effectively support the growth of various strains, but its promotion on protein yield was weak. After optimization, the protein productivity of 0.17 g/L/d obtained in this study was only lower than the data reported by Patel, Krikigianni, Rova, Christakopoulos and Matsakas [65] and Wang, Wang, Ding, Yu, Wang, Geng, Li and Wen [66]. The higher tolerance to acetate of those strains definitely made a difference, while the added 30 g/L NaAc (equivalent to 732 mM NaAc-C) caused the complete inhibition of the growth of *C. sorokiniana UTEX 1230*. Additionally, the dark environment appears to favor protein synthesis metabolism [67]. Further, with a well-designed cultivation strategy discussed previously, it is promising to achieve protein yields of 0.34 g/L/d, ranking the highest and thus highlighting the potential of microalgae cultured in this study as a protein source.

**Table 5.**
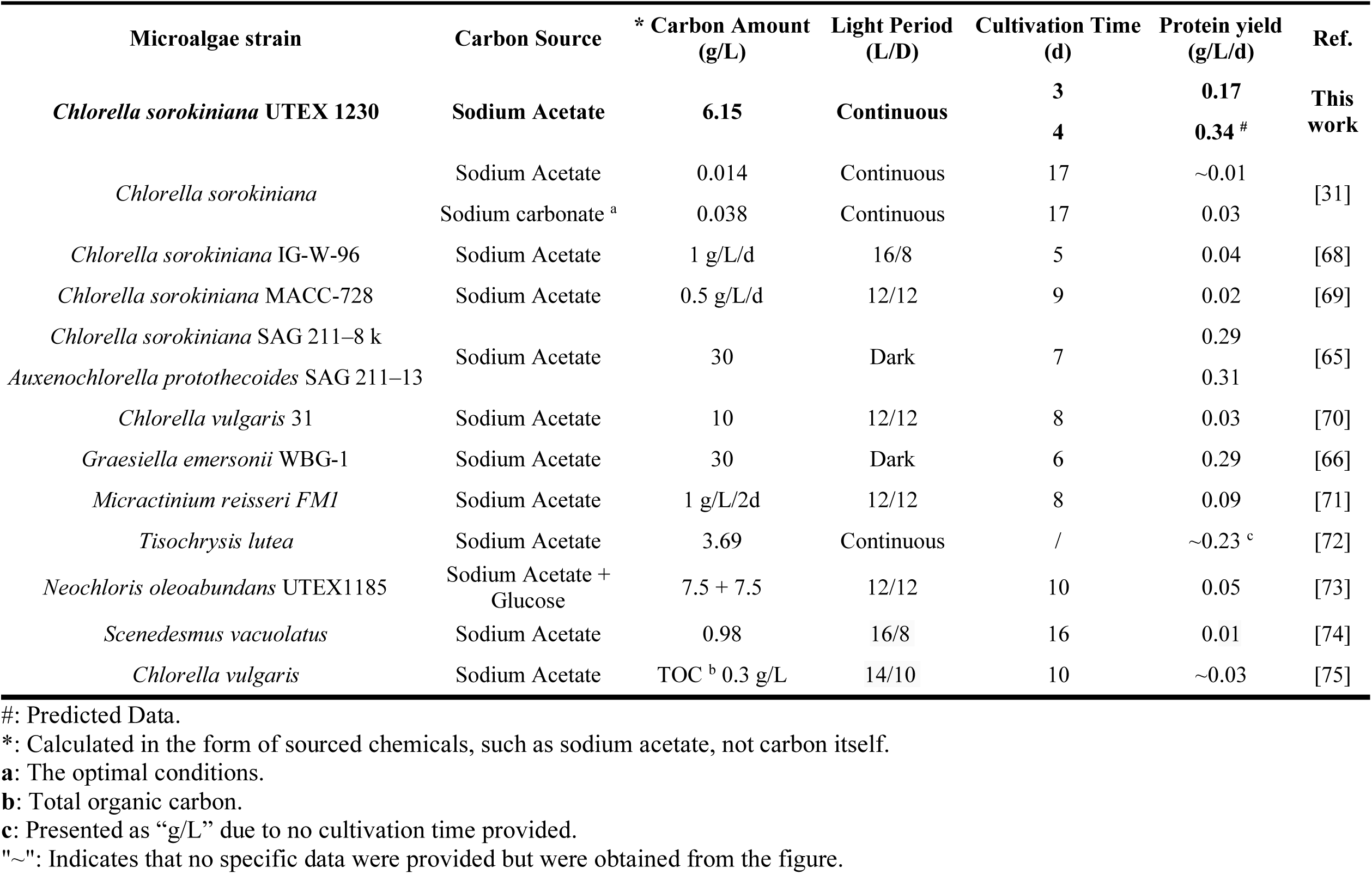
Comparison of Protein Productivity of Microalgae Growing on Acetate according to literature and this work.

## 4. Conclusions

By confirming the ability of *C. sorokiniana* UTEX 1230 to utilize NaAc-C compared to glucose, and by optimizing initial conditions such as an inoculum size with an OD of 0.1 and the introduction of 5% CO_2_ for shorter cultivation periods and stable pH levels, this study paves the way for examining how NaAc-C and nitrate-sourced nitrogen inputs impacted the microalgal growth profile. Definitely, the optimal NaAc-C (150 mM) and nitrogen (750 mg/L as NaNO_3_) combination strongly favored the mixotrophic growth of the microalgae. Monitoring carbon and nitrogen consumption clarified variations in biomass accumulation and protein synthesis, with detailed analysis of protein changes over time informing the cultivation and collection strategy for high protein productivity. This study is the first work to unravel the biomass production, especially protein, of *C. sorokiniana* UTEX 1230 under the mixotrophic mode with acetate being the only organic carbon source, but the physicochemical and functional properties of this microalgal protein remain to be determined in the future. Besides, in-depth research is warranted to explore fed-batch or continuous cultivation to address the trade-off between high biomass concentration and extended lag phases due to high NaAc-C and nitrogen feeding, thus achieving the continuous harvest of high-valued biomass. Overall, this research method serves as a guide for investigating other microalgal strains for improved biomass quality and provides theoretical insights for refining the microalgae as an alternative protein or other food additives.

## Data availability

Data will be made available on request.

## Acknowledgments

The project was financially supported by National Science Foundation Future Manufacturing Seed Grant with award # 2328159, and United States Department of Agriculture Sun Grant Program Northeast Region with award # 240583. We extend our special thanks to Honglin Zhu for technical assistance with TGA and N elemental analysis, and to Tiangang Yang and Dr. Jie He for their expertise and support with HPLC technology.

