## Supplemental Materials for "Impact of Acetate and Optimized Nitrate Levels on Mixotrophic Growth and Protein Dynamics in *Chlorella Sorokiniana*"

| Table S1. Formula of Bold's Basal Medium (mg/L) | |
|---|---|
| Boric Acid | 11.42 |
| Calcium Chloride, Anhydrous | 18.872 |
| Cobalt Nitrate•6H2O | 0.49 |
| Cupric Sulfate•5H2O | 1.57 |
| EDTA, Disodium Salt | 63.61 |
| Ferrous Sulfate•7H2O | 4.98 |
| Magnesium Sulfate, Anhydrous | 36.626 |
| Zinc Sulfate•7H2O | 8.82 |
| Manganese Chloride•4H2O | 1.44 |
| Sodium Molybdate | 1.194 |
| Potassium Hydroxide | 31 |
| Potassium Phosphate, Dibasic | 75 |
| Potassium Phosphate, Monobasic | 175 |
| Sodium Chloride | 25 |
| Sodium Nitrate | 250 |

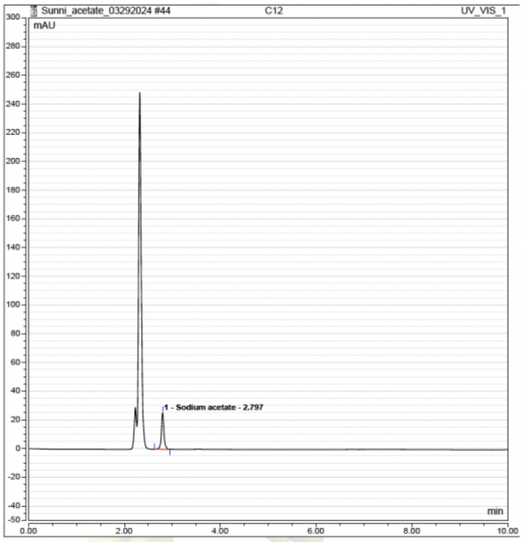


**Figure S1**. HPLC chromatogram of NaAc.

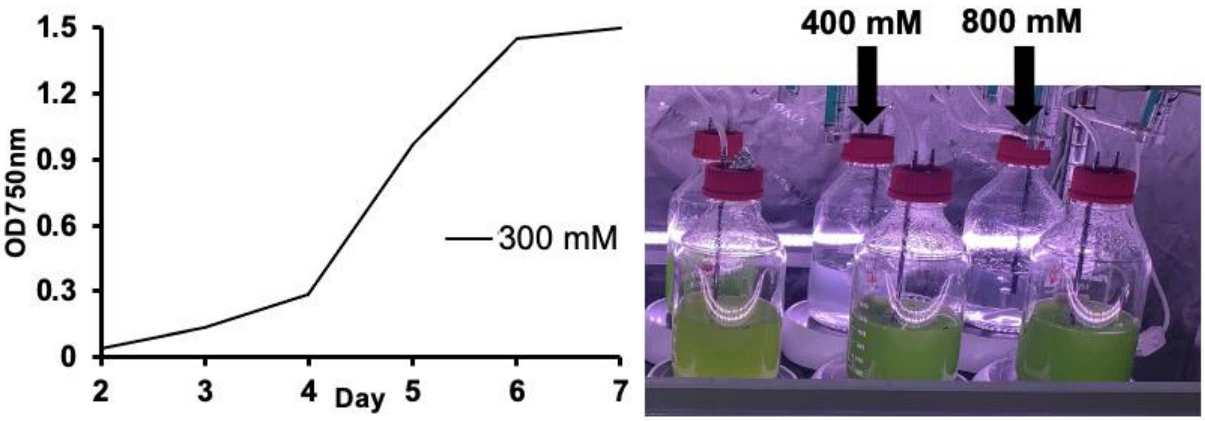


**Figure S2**. Effect of NaAc-C concentration on the growth of microalgae, data from Run #1.

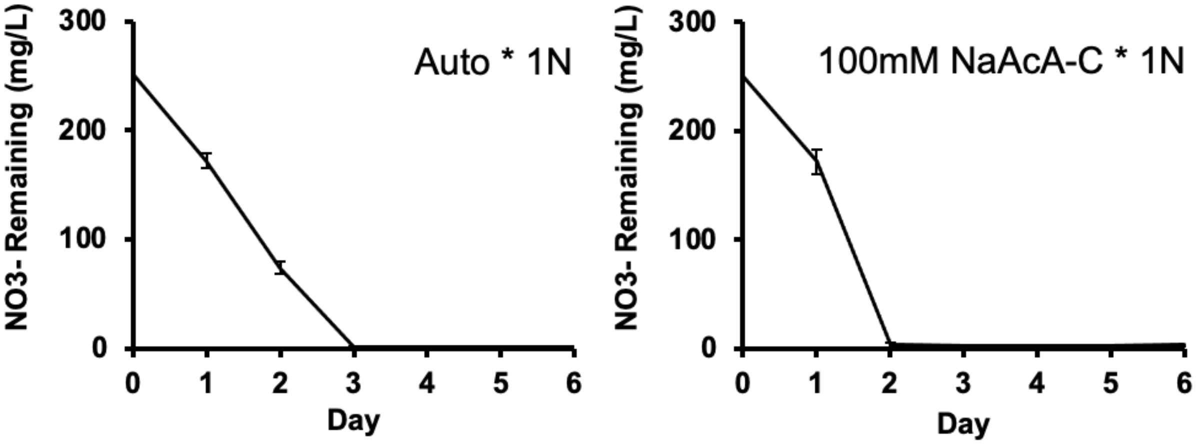


**Figure S3**. The $NO_3^-$ remaining under autotrophic and mixotrophic cultivation (100 mM NaAc-C). 1N was 250 mg/L $NaNO_3$.

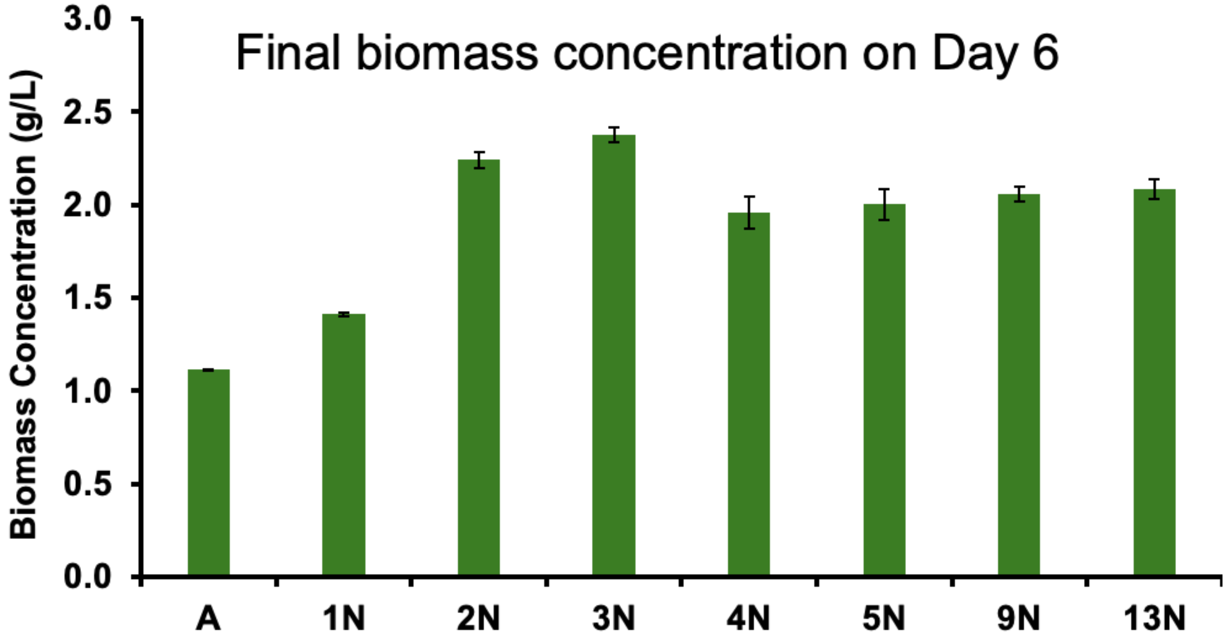


**Figure S4**. Biomass concentration under different $NO_3^-$ levels with 100 mM NaAc-C, data from Run #1. 1N was 250 mg/L $NaNO_3$, 2N was 500 mg/L, and so forth.

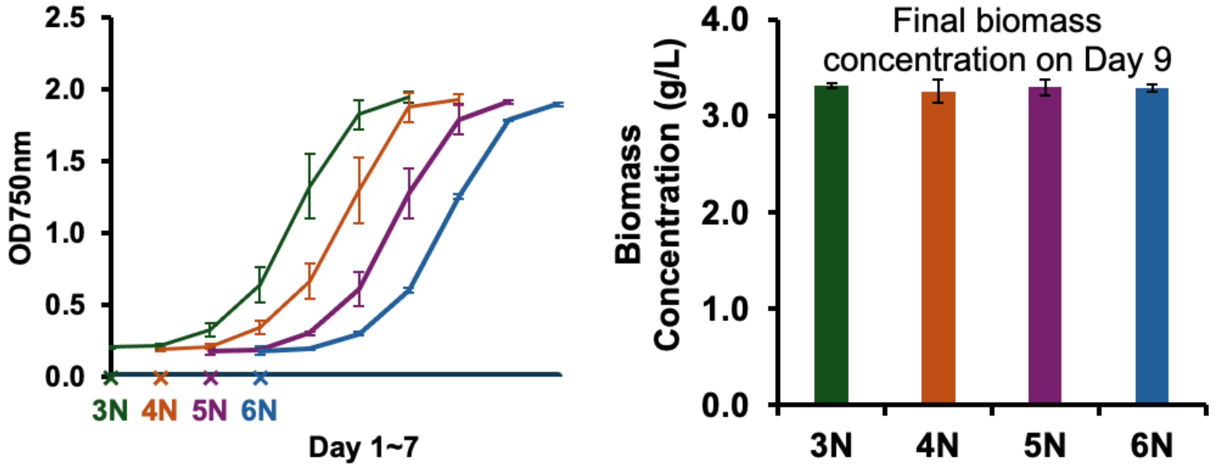


**Figure S5**. Effect of $NaNO_3$ concentrations on (left) growth curve treated with 200 mM NaAc-C and (right) the final biomass concentration collected on the $9^{th}$ day. 1N was 250 mg/L $NaNO_3$, 2N was 500 mg/L, and so forth.

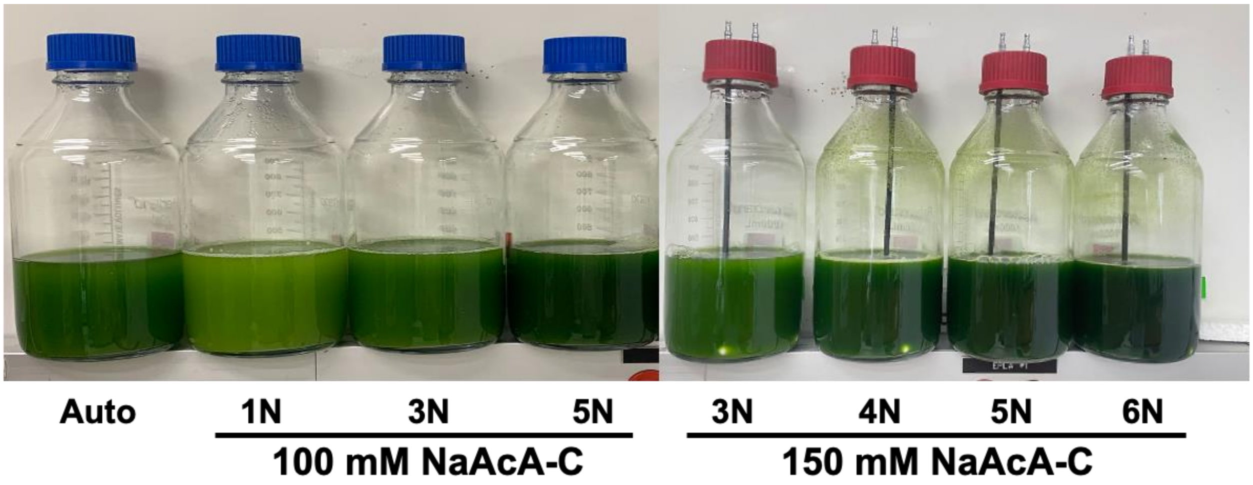


**Figure S6**. Picture of microalgal suspension with different nitrogen levels, supplemented with 100 mM NaAc-C (blue cap) or 150 mM NaAc-C (red cap), on the collection day (6$^{th}$). 1N was 250 mg/L $NaNO_3$, 2N was 500 mg/L, and so forth.
